## Supplementary Appendix for "Spatial structure facilitates evolutionary rescue by drug resistance"

**4 Lattice, star and line structures with sensitive inoculum** **13**

**5 Clique population structure with mutants in the inoculum** **15**

**6 Concrete examples of spatially structured populations** **16**

#### 1 Derivation of analytical results presented in the main text

We describe population composition using a stochastic model (see "Models and methods" in the main text). This implies that the number $N$ of bacteria in a deme is constantly fluctuating in time. However, because $K \gg 1$ and $g \ll f_S = 1$, these fluctuations around the steady-state deme size $N^*$ remain small. At steady state, the division rate balances the death rate, leading to: $f_S(1-N^*/K) = g$ in a population that only comprises S bacteria. This yields

$$N^* = K\left(1 - g/f_S\right) \tag{S1}$$

In all the analytical calculations that follow, we approximate deme size $N$ by $N^*$ (see Ref. [1]).

#### 1.1 Probability of presence of mutants from a lineage destined to go extinct

Let us consider a deme where R mutants have not fixed, in the absence of drug. Let us determine the probability $p_{\mathrm{pres}}$ of presence of mutants that arise through mutations upon division of sensitive individuals, and whose lineage is destined to go extinct in the absence of drug, but does not undergo rapid stochastic extinction after drug is added. We follow the method exposed in Ref. [1], which is based on the transition rate matrix of the process.

**Presence of $i$ mutants from a lineage destined to go extinct without drug.** To incorporate the stochastic extinction effect, we need to distinguish each case where exactly $i$ mutants are present, as the probability of stochastic extinction will depend on $i$. The probability of having exactly $i$ mutants present in a well-mixed population with $N^*$ individuals reads $p_R(i) = \tau_R^d(i)\, N^*\mu g$, where $\tau_R^d(i)$ is the average time spent in a state with $i$ mutants by a resistant lineage destined for extinction (known as the sojourn time [2]), and $N^*\mu g$ the total mutation rate, with $g$ the death rate and $\mu$ the mutation probability upon division. We have $\tau_R^d(i) = -\pi_i/\pi_1(\tilde{\mathbf{R}}^{-1})_{i1}$, where $\pi_i$ is the probability that resistant mutants go extinct, starting from $i$ of them, while $\tilde{\mathbf{R}}$ is the reduced transition rate matrix for mutants (i.e., the transition rate matrix where rows and columns corresponding to absorbing states are eliminated), see Ref. [1]. This yields the probability that $i$ mutants from a lineage destined to go extinct are present in the deme:

$$p_R(i) = -N^* g\mu \,\frac{\pi_i}{\pi_1}(\tilde{\mathbf{R}}^{-1})_{i1}\,. \tag{S2}$$

As discussed in the main text, when drug is added to the system, the population can survive if mutants are present when the drug is added. For survival, it is also necessary that the mutant lineages do not quickly go stochastically extinct after drug is added. Let us thus calculate the probability that such a stochastic extinction occurs.

**Probability of stochastic extinction.** Let us consider one resistant bacterium with replication rate $f_R$ and death rate $g$. Let us denote by $e$ the extinction probability of its lineage. This resistant individual can die without dividing with probability $p_D = g/(f_R+g)$, giving an extinction probability of $e_D = 1$. It can also divide before dying with probability $p_R = f_R/(f_R+g)$, leading to the extinction probability $e_R = e^2$, assuming independence of lineages, in the branching process framework [3]. We can then write an equation for the extinction probability $e$ of the lineage of the single resistant bacterium:

$$e = p_D\, e_D + p_R\, e_R = \frac{g}{f_R+g} + e^2 \frac{f_R}{f_R+g}. \tag{S3}$$

The solutions to this equation are $e = 1$ or $e = g/f_R$ if $g < f_R$. Thus, assuming again that the fate of each lineage is independent, we obtain an extinction probability $(g/f_R)^i$ starting from $i$ mutants.

**Presence of a mutant lineage destined to go extinct without drug, but that does not undergo stochastic extinction when drug is added.** We finally combine Eq S2 with the stochastic extinction probability to find:

$$p_{\mathrm{pres}} = \sum_{i=1}^{N^*} \left[1-(g/f_R)^i\right] p_R(i) = -\sum_{i=1}^{N^*} \left[1-(g/f_R)^i\right]\, N^* g\mu \,\frac{\pi_i}{\pi_1}(\tilde{\mathbf{R}}^{-1})_{i1}\,. \tag{S4}$$

Eq S4 is a good approximation of the survival probability of the population on timescales smaller than the average time of appearance of a successful R mutant, since R mutants that appear before are doomed to go extinct in the absence of drug.

**Lifetime of a mutant lineage destined to go extinct without drug.** In the absence of drug, the total average lifetime $\tau_R^d$ of a mutant lineage destined to go extinct can be obtained by summing the sojourn times $\tau_R^d(i)$ expressed above (see also Ref. [2]):

$$\tau_R^d = \sum_{i=1}^{N^*} \tau_R^d(i) = -\sum_{i=1}^{N^*} \frac{\pi_i}{\pi_1}(\tilde{\mathbf{R}}^{-1})_{i1}\,. \tag{S5}$$

For neutral R mutants, this gives

$$\tau_R^d = \frac{1}{g}\left(\frac{N^*}{N^*-1}\sum_{i=1}^{N^*-1}\frac{1}{i} - 1\right) \approx \frac{1}{g}\log(N^*)\,, \tag{S6}$$

where we assumed $N^* \gg 1$ to obtain the last expression. This lifetime is only weakly impacted by population size.

#### 1.2 Appearance of a mutant that fixes in a well-mixed population

In our model, mutations from sensitive to resistant individuals happen at birth with probability $\mu$. Since at equilibrium $f_S(1 - N^*/K) = g$, the average time it takes for a mutant to appear in a population of $N^*$ wild-types is in the rare mutation regime:

$$\langle t_{\mathrm{app}}(N^*)\rangle = \frac{1}{N^*\mu g}. \tag{S7}$$

Thus, the average time $\langle t_{af\,W}(N^*)\rangle$ of appearance of a mutant that fixes in a well-mixed population of fixed size $N^*$ is given by:

$$\langle t_{af\,W}(N^*)\rangle = \langle t_{\mathrm{app}}(N^*)\rangle \times \frac{1}{p_{\mathrm{fix}}(N^*)} = \frac{1}{N^*\mu g\, p_{\mathrm{fix}}(N^*)}. \tag{S8}$$

**Cost-free mutants.** The fixation probability of neutral mutants in a population of size $N^*$ is:

$$p_{\mathrm{fix}}(N^*) = \frac{1}{N^*}\,. \tag{S9}$$

Thus, the average time $\langle t_{af\,W}\rangle$ of appearance of a neutral mutant that fixes in a well-mixed population of fixed size $N^*$ is given by:

$$\langle t_{af\,W}(N^*)\rangle = \langle t_{\mathrm{app}}(N^*)\rangle \times \frac{1}{p_{\mathrm{fix}}(N^*)} = \frac{1}{\mu g}\,. \tag{S10}$$

This implies that for cost-free mutants, the average time of appearance of a successful mutant is independent of the population size: it remains the same for a single deme and a well-mixed population.

**Mutants with cost $\delta$.** More generally, the fixation probability of a mutant with fitness cost $\delta$ in a well-mixed population of fixed size $N^*$ described by the Moran process is [2]

$$p_{\mathrm{fix}}(N^*) = \frac{(1-\delta)^{-1} - 1}{(1-\delta)^{-N^*} - 1}\,. \tag{S11}$$

If $\delta \ll 1/N^*$, then to leading order

$$p_{\mathrm{fix}}(N^*) = \frac{1}{N^*}\,. \tag{S12}$$

The mutant is then said to be effectively neutral. Conversely, if $\delta \gg 1/N^*$, then to leading order

$$p_{\mathrm{fix}}(N^*) = \delta e^{-N^*\delta}\,. \tag{S13}$$

Mutants with such a substantial fitness cost have a fixation probability that is exponentially suppressed.

In Fig 2B, as $K\delta = 1$, we employ the most general formula for $p_{\mathrm{fix}}$, see Eq S11. The cost regimes leading to the simplified expressions of $p_{\mathrm{fix}}$ in Eqs S12 and S13 are further discussed in Section 3.2.

#### 1.3 Appearance of the slowest mutant that fixes in a structured population

Let us now focus on the appearance of a mutant that fixes in the slowest deme in a population composed of $D$ independent demes which each have steady-state size $N^*$, in the absence of migrations. By *slowest deme*, we mean the deme in which a locally successful mutant (i.e. a mutant destined to fix locally in its deme of origin) takes the longest time to appear.

For a mutant to fix in the slowest deme, mutants must have fixed in all others. Let us denote by $t_{af\,S}$ (resp. $t_{af\,d}$) the time of appearance of a mutant destined to fix in the slowest deme (resp. in any deme). The probability $P(t_{af\,S} \leq t)$ that $t_{af\,S}$ is smaller or equal than $t$ reads for any $t$:

$$P(t_{af\,S} \leq t) = [P(t_{af\,d} \leq t)]^D . \tag{S14}$$

The appearance of a locally successful mutant is a Poisson process with rate $\lambda = N^* \mu g\, p_{\text{fix}}(N^*)$ (corresponding to the inverse of Eq S8). Thus, the time $t_{af\,d}$ is exponentially distributed with rate $\lambda$ and probability density

$$p_d(t_{af\,d} = t) = \lambda e^{-\lambda t} , \tag{S15}$$

and we have $P(t_{af\,d} \leq t) = 1 - e^{-\lambda t}$. This allows us to express the probability density $p_S(t)$ of appearance of a mutant in the slowest deme:

$$\begin{aligned} p_S(t)dt &= dp(t_{af\,S} \in [t, t+dt]) = P(t_{af\,S} \leq t+dt) - P(t_{af\,S} \leq t) \\ &= \frac{dP(t_{af\,S} \leq t)}{dt} dt = D \left[P(t_{af\,d} \leq t)\right]^{D-1} \frac{dP(t_{af\,d} \leq t)}{dt} dt \\ &= D \left[1 - e^{-\lambda t}\right]^{D-1} \lambda e^{-\lambda t} dt . \end{aligned} \tag{S16}$$

Note that $p_S(t)$ is positive for all $t$ and normalized, as expected.

The average appearance time of a locally successful mutant in the slowest deme is thus given by:

$$\langle t_{af\,S}(N^*, D) \rangle = \int_0^\infty t\, p_S(t)\, dt = \frac{1}{\lambda} \sum_{i=1}^{D} \frac{1}{i} , \tag{S17}$$

which involves the harmonic number $\sum_{i=1}^{D} 1/i$.

#### 1.4 Appearance of the fastest mutant that fixes in a structured population

Let us now consider the *fastest deme*, i.e. the deme where a locally successful mutant first appears. By definition, if a successful mutant takes longer than $t$ to appear in the fastest deme, it entails that successful mutants will take longer to appear in all demes. Denoting by $t_{af\,F}$ the time of appearance of a successful mutant in the fastest deme, we have:

$$P(t_{af\,F} > t) = [P(t_{af\,d} > t)]^D . \tag{S18}$$

This allows us to express the probability density $p_F$ of appearance of a successful mutant in the fastest deme:

$$\begin{aligned} p_F(t)dt &= dp(t_{af\,F} \in [t, t+dt]) = -\frac{dP(t_{af\,F} > t)}{dt} dt \\ &= -D[P(t_{af\,d} > t)]^{D-1} \frac{dP(t_{af\,d} > t)}{dt} dt \\ &= -D[1 - P(t_{af\,d} \leq t)]^{D-1} \left[-\frac{dP(t_{af\,d} \leq t)}{dt}\right] dt \\ &= D\lambda e^{-\lambda D t} dt , \end{aligned} \tag{S19}$$

with $\lambda = N^* \mu g\, p_{\text{fix}}(N^*)$, as previously defined. Note that $p_F(t)$ is positive for all $t$ and normalized, as expected.

The average appearance time of a successful mutant in the fastest deme is thus given by (see also Ref. [4]):

$$\langle t_{af\,F}(N^*, D) \rangle = \int_0^\infty t\, p_F(t)\, dt = \frac{1}{D\lambda} . \tag{S20}$$

In the case of cost-free mutants, using Eq S9 gives

$$\langle t_{af\,F}(N^*, D) \rangle = \frac{1}{D\mu g} , \tag{S21}$$

In the case where resistance carries a cost, using Eq S11 gives

$$\langle t_{af\,F}(N^*, D) \rangle = \frac{1}{DN^* \mu g} \frac{(1-\delta)^{-N^*} - 1}{(1-\delta)^{-1} - 1} . \tag{S22}$$

#### 1.5 Survival probability of the population with $\gamma = 0$

When there is no migration between demes, i.e. when $\gamma = 0$, the survival probability $p_{s\,\gamma=0}(D)$ of a population comprising $D$ demes, is related to the survival probability $p_{s\,d}$ of a deme through:

$$p_{s\,\gamma=0}(D) = 1 - (1 - p_{s\,d})^D. \tag{S23}$$

Indeed, denoting by $p_{e\,\gamma=0}$ the probability of extinction of all demes, we have:

$$p_{s\,\gamma=0}(D) = 1 - p_{e\,\gamma=0}(D). \tag{S24}$$

Let $p_{e\,d}$ be the probability that the population in one deme becomes extinct. When $\gamma = 0$, extinctions in different demes are independent events with the same probability, thus:

$$p_{s\,\gamma=0}(D) = 1 - p_{e\,\gamma=0}(D) = 1 - (p_{e\,d})^D = 1 - (1 - p_{s\,d})^D. \tag{S25}$$

We checked that Eq S23 was satisfied in our numerical simulations, within the errors given by the standard error of the proportion.

#### 1.6 Closed form of the survival probability for well-mixed and fully subdivided populations

In our structured populations, mutants appear at random in any deme. The population can survive in two distinct ways:

- Due to the presence of successful lineages that have fixed locally or globally in the population, and allow survival when drug is added.
- Due to the presence of a mutant lineage destined for extinction in the absence of drug at the time $t_{\text{add}}$ when drug is added. This mutant lineage can proliferate in the presence of drug. The population then survives, unless this mutant lineage undergoes a quick stochastic extinction.

The survival probability can be then written by summing over these two distinct scenarios:

$$\begin{aligned} p_{\text{surv}}(t_{\text{add}}) &= p_{\text{succ, surv}}(t_{\text{add}}) + p_{\text{no succ, surv}}(t_{\text{add}}) \\ &= p_{\text{succ}}(t_{\text{add}})\, p_{\text{surv}|\text{succ}} + [1 - p_{\text{succ}}(t_{\text{add}})]\, p_{\text{surv}|\text{no succ}} \\ &= p_{\text{succ}}(t_{\text{add}}) + [1 - p_{\text{succ}}(t_{\text{add}})]\, p_{\text{pres}}. \end{aligned} \tag{S26}$$

In the first line, we denoted by $p_{\text{succ, surv}}(t_{\text{add}})$ the probability that a successful mutant has appeared by $t_{\text{add}}$ and leads to population survival, and by $p_{\text{no succ, surv}}(t_{\text{add}})$ the probability that no successful mutant has appeared at $t_{\text{add}}$ but that the population survives (thanks to the presence of a mutant lineage that was destined for extinction). In the second line, $p_{\text{succ}}(t_{\text{add}})$ is the probability that a successful mutant has appeared by $t_{\text{add}}$. The probability $p_{\text{surv}|\text{succ}}$ of survival conditioned on the presence of a successful mutant is one, while the probability $p_{\text{surv}|\text{no succ}}$ of survival conditioned on the absence of any successful mutant is $p_{\text{pres}}$, given by Eq S4.

Let us now calculate $p_{\text{succ}}(t_{\text{add}})$ in a well-mixed population of fixed size $N^*$. Given that the appearance of a successful mutant is a Poisson process with rate $\lambda = N^* \mu g p_{\text{fix}}(N^*)$, the associated time of appearance is distributed according to Eq S15. Thus, $p_{\text{succ}}(t_{\text{add}})$ reads:

$$p_{\text{succ}}(t_{\text{add}}) = \int_0^{t_{\text{add}}} \lambda e^{-\lambda t} dt = 1 - e^{-\lambda t_{\text{add}}}. \tag{S27}$$

For neutral mutants in a well-mixed population of size $N^*$, this reads:

$$p_{\text{succ}}(t_{\text{add}}) = 1 - e^{-\mu g t_{\text{add}}}, \tag{S28}$$

while for deleterious mutants in a well-mixed population of size $N^*$ we have:

$$p_{\text{succ}}(t_{\text{add}}) = 1 - \exp\left(-N^* \mu g \frac{(1-\delta)^{-1} - 1}{(1-\delta)^{-N^*} - 1} t_{\text{add}}\right). \tag{S29}$$

Let us now consider a structured population, and ask whether a locally successful mutant has appeared by $t_{\text{add}}$. For the appearance time of successful mutants in the fastest deme of a structured population, the probability density is given by Eq S19, yielding:

$$p_{\text{succ}}(t_{\text{add}}) = \int_0^{t_{\text{add}}} D\lambda e^{-\lambda D t} dt = 1 - e^{D\lambda t_{\text{add}}} . \tag{S30}$$

If mutants are neutral, we have:

$$p_{\text{succ}}(t_{\text{add}}) = 1 - e^{-D\mu g t_{\text{add}}} , \tag{S31}$$

and if they carry a resistance cost:

$$p_{\text{succ}}(t_{\text{add}}) = 1 - \exp\left(-DN^*\mu g \frac{(1-\delta)^{-1} - 1}{(1-\delta)^{-N*} - 1} t_{\text{add}}\right) . \tag{S32}$$

### 2 Description of the simulations performed in the main text

#### 2.1 General simulation approach

Our simulations are based on the Gillespie algorithm [5–7]. Let us consider a clique population structure composed of $D$ demes, labeled with $i \in [1, D]$. Let $S_i$ (resp. $R_i$) denote one sensitive (resp. resistant) individual in deme $i$. The system obeys the following reaction network:

$$\begin{aligned}
&S_i \xrightarrow{f_S\left(1-\frac{N}{K}\right)(1-\mu)} 2S_i \, , \\
&S_i \xrightarrow{f_S\left(1-\frac{N}{K}\right)\mu} S_i + R_i \, , \\
&S_i \xrightarrow{g} \emptyset \, , \\
&S_i \xrightarrow{\gamma} S_j \;\text{ for all }\; j \neq i \, , \\
&R_i \xrightarrow{f_R\left(1-\frac{N}{K}\right)} 2R_i \, , \\
&R_i \xrightarrow{g} \emptyset \, , \\
&R_i \xrightarrow{\gamma} R_j \;\text{ for all }\; j \neq i \, .
\end{aligned} \tag{S33}$$

The action of the biostatic drug is modeled through a change of the fitness of sensitive individuals, $f_S$, as follows:

$$f_S = \begin{cases} 1 & \text{in the absence of drug,} \\ 0 & \text{after biostatic drug addition.} \end{cases} \tag{S34}$$

**Time until colonization of next deme.** In Fig 5A, we perform simulations to assess the time until colonization of next deme. These simulation are set up slightly differently than others, to focus on the colonization process. We initialize one of the demes with $N = 0.9 \times K$ mutants, all others with $N = 0.9 \times K$ wild-types, mimicking fixation of mutants in one deme before drug addition. We add the biostatic drug at $t = 0$ in this simulation, thus modeling the case where one deme has fixed resistance before drug addition. We then let the system evolve until overall colonization of the structured population by resistant mutants. For each $k \geq 1$, we save the times of successful colonization of the $(k+1)^{th}$ deme, given that $k$ demes were already colonized by mutants. We define successful colonization of a deme as reaching a number of mutants $0.9 \times K$. We then obtain the values of $\langle t_{\mathrm{c\ mig}}(k) \rangle$ as the differences between the successive times we recorded.

**Time until overall colonization.** In Fig 5B, we report the time until overall colonization of the structured population by resistant mutants. When saving the state of the system after overall mutant colonization (resp. extinction), we also record the time it took for overall colonization (resp. extinction). Thus, we can calculate the mean of the overall colonization time for each $t_{\mathrm{add}}$ and each structure. Note that the fixation of resistance may happen before drug addition (this happens in particular for addition times longer than $\langle t_{af\,S} \rangle$). In this case, we consider that colonization happens when resistant individuals reach fixation in all demes and there are at least $0.9 \times K$ resistant individuals.

### 3 Clique population structure with sensitive inoculum

#### 3.1 Clique population structure with different numbers of demes

To assess the impact of the degree of subdivision of a population on rescue by resistance, we extend the study of the survival probability with neutral R mutants in Fig 2A to different numbers of demes. Fig S1 shows the impact of varying the degree of subdivision of a population of fixed total size $DK$, but composed of a different number of demes ($D = 5, 10, 20$). We observe that higher subdivision yields higher probabilities of survival.

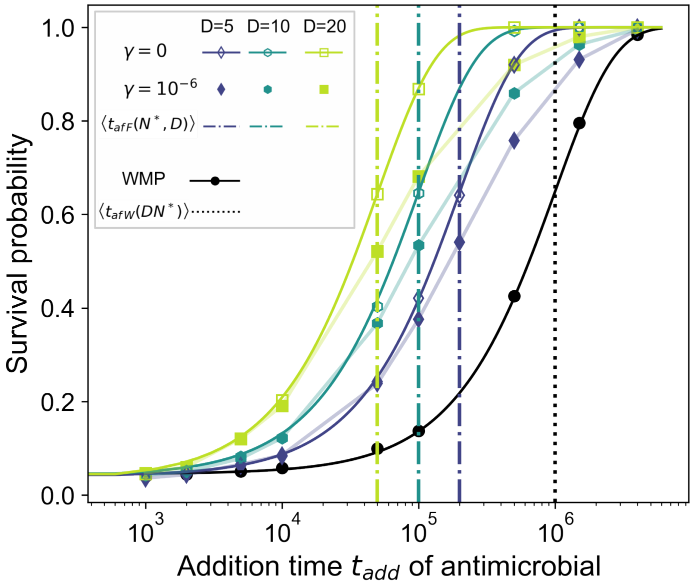

Figure S1: **Survival probability of a bacterial population with neutral mutants for varying degree of subdivision at a fixed total population size.** Survival probability as a function of the treatment addition time $t_{\text{add}}$. Three levels of subdivision of a population with total carrying capacity $DK = 1000$, into 5, 10, and 20 demes, are shown for two values of $\gamma$. Results for the well-mixed population ("WMP") are shown for reference. In all cases, we determined the survival probability from the fraction of $10^4$ simulations where the population survived after drug application. The vertical dash-dotted lines denote the average time of appearance of a locally successful mutant in the fastest deme of each structure (Eq 2). The vertical dotted line represents the average appearance time of a successful mutant in a well-mixed population (Eq 1). Solid lines for the fully subdivided ($\gamma = 0$) and well-mixed populations are analytical predictions from Eq 4. Thin lines connecting data points are guides for the eye. Parameter values: $f_S = 1$ without drug, $f_S = 0$ with drug, $f_R = 1$ (no cost of resistance), $g = 0.1$, $\mu = 10^{-5}$.

#### 3.2 Impact of a cost or of a benefit of resistance

In Fig 2, we considered the case where the resistant mutant is neutral in the absence of drug and the case where it carries a moderate cost of resistance $\delta = 1/K$. Here, we discuss different regimes of cost of resistance, and present simulation results for a higher cost of resistance. Besides, if antibiotic is already present at relatively low doses in the environment before the addition of drug, it can induce a benefit of resistance before drug is added. We briefly discuss this case at the end of this section.

**Different regimes of cost.** Let us consider different regimes of cost for R mutants.

As discussed in the main text, for neutral resistant mutants ($\delta = 0$), $p_{\text{fix}} = 1/N^*$, so Eq 1 gives $\langle t_{af\,W}(N^*)\rangle = 1/(\mu g)$. Importantly, this result does not depend on population size $N^*$. Therefore, it holds both for each deme in a structured population and for a well-mixed population of steady-state size $DN^*$: $\langle t_{af\,W}(DN^*)\rangle = 1/(\mu g)$. Meanwhile, Eq 2 gives $\langle t_{af\,F}(N^*, D)\rangle = 1/(D\mu g) = \langle t_{af\,W}(N^*)\rangle/D$. Therefore, a locally successful mutant appears in a structured population $D$ times faster than a successful mutant in a well-mixed population with same total steady-state size $DN^*$.

For mutants with a substantial cost of resistance $\delta \gg 1/K$, $p_{\text{fix}} = \delta e^{-N^*\delta}$ (see Eq S13), and thus $\langle t_{af\,W}(N^*)\rangle = e^{N^*\delta}/(N^*\mu g\delta)$, which is exponentially longer than in the neutral case and satisfies $\langle t_{af\,W}(N^*)\rangle \ll \langle t_{af\,W}(DN^*)\rangle$. Therefore, $\langle t_{af\,F}(N^*, D)\rangle = \langle t_{af\,W}(N^*)\rangle/D \ll \langle t_{af\,W}(DN^*)\rangle$.

Finally, mutants with intermediate cost satisfying $1/(DK) \ll \delta \ll 1/K$ are effectively neutral in demes of size $N^*$, leading to $\langle t_{af\,F}(N^*, D)\rangle = 1/(D\mu g)$ as in the neutral case (see Eq S12). However, they are substantially deleterious in a well-mixed population of size $DN^*$, leading to $\langle t_{af\,W}(DN^*)\rangle = e^{DN^*\delta}/(DN^*\mu g\delta)$, which is exponentially larger than $\langle t_{af\,F}(N^*, D)\rangle$.

**Strong cost of resistance.** With a strong cost of resistance $\delta$ satisfying $\delta \ll 1$ and $N^*\delta \gg 1$ (satisfied in Fig S2), the probability of fixation of one mutant in a deme of size $N^*$ is approximately $\delta e^{-N^*\delta}$ (cf. Eq S13). It is thus exponentially suppressed, and the time to appearance of a successful mutant in a well-mixed population of size $N^*$ is exponentially longer than in the neutral case. With the parameter values of Fig S2, the appearance time of a successful mutant in a well-mixed population of size $DN^*$ is $\langle t_{af\,W}(DN^*)\rangle \approx 1.3 \times 10^{43}$, while the appearance time of the fastest locally successful mutant in a population of $D$ demes of size $N^*$ each is $\langle t_{af\,F}(N^*, D)\rangle \approx 1.3 \times 10^8$. Therefore, fixation events can be neglected in the range of $t_{\mathrm{add}}$ considered previously for neutral mutants and mutants with moderate cost (see Fig 2 for the clique, and in Fig S6 for the star). Thus, in this range of $t_{\mathrm{add}}$, we can estimate the survival probability from the probability of presence of mutant lineages destined for extinction, introduced in Eq S4. This analytical prediction agrees with our simulation results, see Fig S2. We do not observe a statistically significant impact of population structure on the survival probability of the population, in this range of $t_{\mathrm{add}}$. Recall however that if times comparable to the mean fixation time of a successful mutant in a deme (or larger) were considered, we predict that spatial structure would favor population survival for strong costs of resistance. We do not consider this further because of the extremely long times involved.

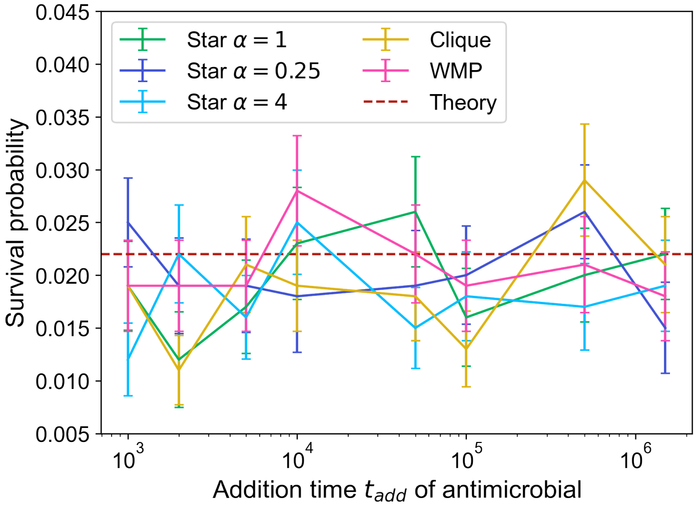

Figure S2: **Survival probability of a structured bacterial population with a cost of resistance.** The survival probability of the population to the addition of biostatic drug is shown versus the drug addition time. We consider a clique and a star with different values of migration asymmetry $\alpha$. The case of the well-mixed population with the same total size ("WMP") is shown for reference. The migration rate is $\gamma = 10^{-6}$ in the clique, while in the star we follow the convention in Section 4 to compare with a clique of a given migration rate $\gamma$, leading to $\gamma_O = (D-1)\gamma$ and $\gamma_I = \alpha\gamma_O$. Simulation results are obtained from $10^3$ replicates, with error bars representing 95% confidence intervals. Red horizontal dashed line ("Theory"): analytical prediction from Eq S4. Parameter values: $K = 200$, $D = 5$, $f_S = 1$ without drug, $f_S = 0$ with drug, $f_R = 0.9$ with and without drug, $g = 0.1$, $\mu = 10^{-5}$.

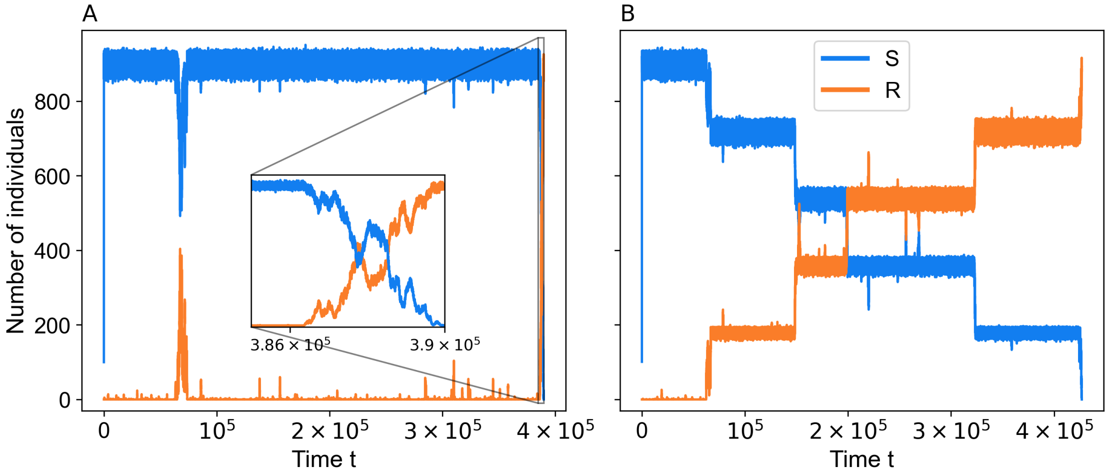

Figure S3: **Dynamics of population in a well-mixed and a structured population.** We present one trajectory from a single simulation realization in the absence of drug for a well-mixed population in Panel A, and one for a structured population with same total size in Panel B. In both cases, we show the number of sensitive (S) and resistant (R) individuals versus time. Parameter values: $K = 200$, $D = 5$, $\mu = 10^{-5}$, $f_S = f_R = 1$ (neutral mutants), $g = 0.1$, $\gamma = 10^{-7}$.

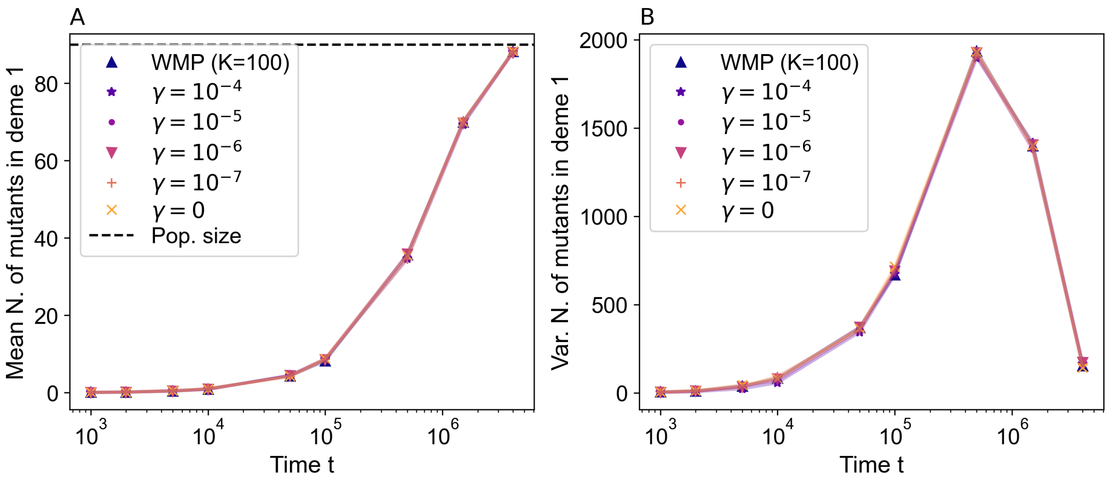

Figure S4: **Mean and variance of the number of mutants at the single-deme level.** Panel A: the mean number of mutants in deme 1 is shown as a function of time in the absence of drug, for different values of the migration rate. Panel B: the variance across simulation replicates of the number of mutants in deme 1 is shown as a function of time in the absence of drug, for different values of the migration rate. Data is obtained from $10^4$ simulation replicates in each case. The case of an isolated deme, i.e. of a well mixed population with carrying capacity $K$ ("WMP ($K = 100$)"), is shown for reference. Parameter values (in both panels): $K = 100$, $D = 10$, $f_S = f_R = 1$ (neutral mutants), $g = 0.1$, $\mu = 10^{-5}$.

##### 3.6 Population composition versus time for different migration rates

In Fig 3C, we analyzed the composition of the population at the time featuring the largest inter-replicate variance for the well-mixed population, namely $t = 5\times10^5$, in spatially structured populations with different migration rates, in the case of neutral R mutants. In Fig S5, we report how this composition evolves in time, in the absence of drug, for spatially structured populations with different migration rates. As in Fig 3C, we focus on the number of mutants in each deme to categorize the population composition.

Recall that here, the population is composed of wild-type sensitive bacteria and of neutral resistant mutants that arise with probability $\mu$ upon division of wild-types. Fig S5 shows that when the per capita migration rate $\gamma$ is decreased, there is a longer phase where the population has an intermediate composition, with some demes harboring a substantial number of resistant mutants ("Big R pop." category). This is consistent with our above discussion in Section 3.3 and with our results in Fig S3. The fixation of mutants occurs gradually, deme by deme, in structured populations with small migration rates. Furthermore, substantial mutant numbers exist earlier in these structured populations. We further observe in Fig S5 that a per capita migration rate $\gamma = 10^{-3}$ results in an evolution of

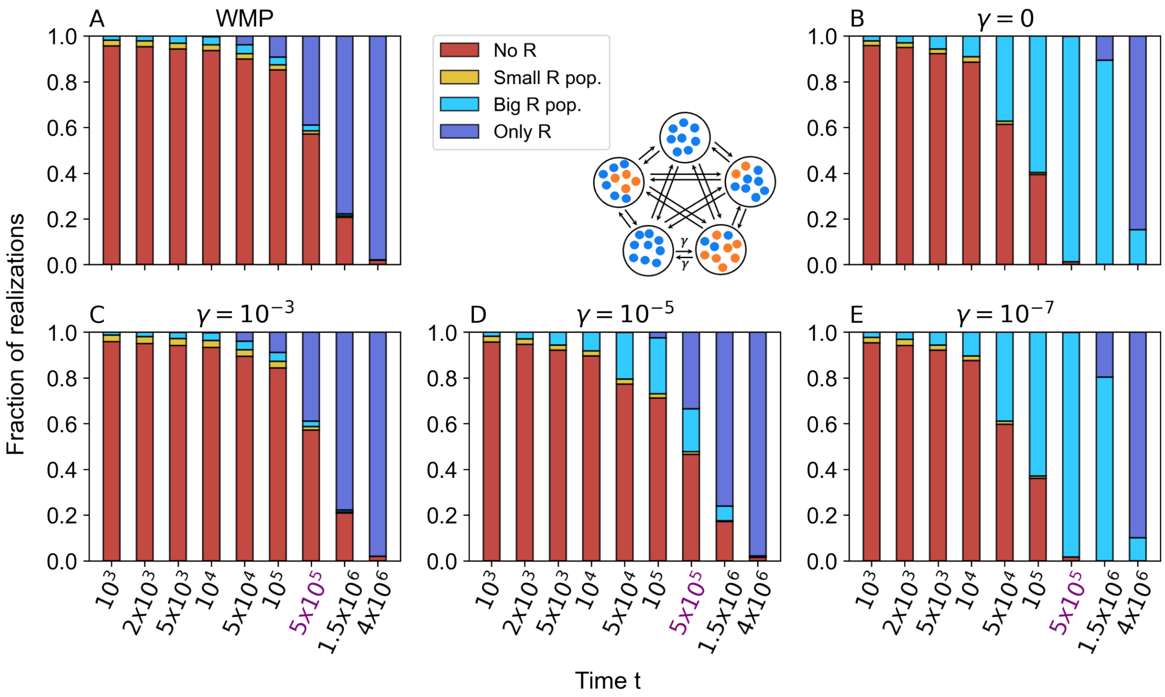

Figure S5: **Population composition versus time for different population structures.** Population composition in the absence of drug in the well-mixed population ("WMP", Panel A), in the fully subdivided population ("SP", Panel B), and in the clique-structured population with same total size for three values of the migration rate $\gamma$ (Panels C-E). The four categories of population composition reported here are based on the number of mutants per deme. They are the same as in Fig 3C, and are defined in the caption of that figure. The x-axis tick corresponding to $t = 5 \times 10^5$, considered in Fig 3C, is highlighted in purple. Data is obtained from $10^4$ simulation replicates in each case. Parameter values (in all panels): $K = 100$, $D = 10$, $f_S = f_R = 1$, $g = 0.1$, $\mu = 10^{-5}$.

population composition akin to that of a well-mixed population, a per capita migration rate $\gamma = 10^{-7}$ yields an evolution of population composition similar to that of a fully subdivided population with $\gamma = 0$. Structures with intermediate values of $\gamma$ feature an intermediate evolution of population composition.

#### 3.7 Colonization timescales after drug is added

In the main text, we discuss the time $\langle t_{\mathrm{c\ mig}} \rangle$ for R mutants to colonize the next empty deme after drug is added, see Eq 7. In Table S1, we report values of $\langle t_{\mathrm{c\ mig}}(k) \rangle$ when $k = 1$ or 5 demes are already mutant, for different values of the migration rate $\gamma$, using the same parameter values as in Fig 5. With these parameters, the decay time upon drug addition of a well-mixed population comprising $N^*$ S bacteria is $\tau_S = 50.8$. Recall that the smallest value of $\langle t_{\mathrm{c\ mig}}(k) \rangle$ is obtained when $k = D/2 = 5$. Thus, the results of Table S1 show that for $\gamma = 10^{-6}$, which is the value used in Fig 5, we have $\tau_S \ll \langle t_{\mathrm{c\ mig}}(k) \rangle$ for all $k$.

| | $\gamma = 10^{-4}$ | $\gamma = 10^{-5}$ | $\gamma = 10^{-6}$ | $\gamma = 10^{-7}$ |
|---|---|---|---|---|
| $\langle t_{\mathrm{c\ mig}}(k = 1) \rangle$ | 13.7 | $1.37 \times 10^2$ | $1.37 \times 10^3$ | $1.37 \times 10^4$ |
| $\langle t_{\mathrm{c\ mig}}(k = 5) \rangle$ | 4.93 | 49.3 | $4.93 \times 10^2$ | $4.93 \times 10^3$ |

Table S1: Numerical evaluation of $\langle t_{\mathrm{c\ mig}} \rangle$ from Eq 7 for $k = 1$ and $k = 5$. Parameter values: $K = 100$, $D = 10$, $g = 0.1$, $f_R = 1$ (neutral mutants), as in Fig 5.

The total colonization time $\langle t_{\text{c tot}} \rangle$ can be obtained by summing $\langle t_{\text{c mig}} \rangle$ over the $D-1$ steps needed to sequentially colonize the structured system. Using Eq 7, we thus obtain:

$$\langle t_{\text{c tot}} \rangle = \sum_{k=1}^{D-1} \langle t_{\text{c mig}}(k) \rangle = \frac{2(\Gamma + \psi(D))}{D\gamma N^*(1 - g/f_R)}\,, \tag{S36}$$

where $\Gamma$ is the Euler gamma constant, while $\psi$ is the digamma function.

### 4 Lattice, star and line structures with sensitive inoculum

Our work mainly focuses on a minimal model of spatially structured populations where all demes are equivalent and connected to one another by identical migration rates. This structure is known as the island model, the clique or the fully connected graph. Let us now extend our study to different graph structures.

**Different structures.** We first consider the grid or square lattice (see Fig S6A). It is a symmetric structure, like the clique, but migrations from each deme are restricted to its four nearest neighbors, thus only allowing local migrations. In the lattice, as in the clique, in each deme the inward migrations balance the outward migrations, i.e. for each $i$ we have $\sum_j \gamma_{ij} = \sum_j \gamma_{ji}$, where $\gamma_{ij}$ denotes the migration rate from deme $i$ to deme $j$: this means that the lattice is a circulation [8–10]. The per capita migration rate in the lattice is denoted by $\gamma_G$, equal in all directions.

We further consider the star (see Fig S6B), comprising a central deme connected to $D-1$ leaves [9]. All leaves are assumed to be equivalent. It is less symmetric than the clique or the square lattice in the sense that the central deme is different from the leaf demes. Migrations from a leaf to the center occur at per capita rate $\gamma_I$, while migrations from the center to the leaf occur at per capita rate $\gamma_O$. We define the migration asymmetry parameter as $\alpha = \gamma_I/\gamma_O$. If $\alpha = 1$, the star is a circulation, while for all other values of $\alpha$ it is not, which impacts the fixation probability of a mutant in the whole population [9, 10].

Finally, we consider a line (see Fig S6C). Migrations from the left to the right occur at per capita rate $\gamma_R$, from the right to the left at rate $\gamma_L$. An asymmetry parameter is defined for the line as well: $\alpha = \gamma_R/\gamma_L$. The structure is symmetric under the simultaneous transformations $\alpha \to 1/\alpha$, $\gamma_R \to \gamma_L$, and $\gamma_L \to \gamma_R$. As for the star, setting $\alpha = 1$ results in a circulation, while $\alpha \neq 1$ impacts the mutant fixation probability [11].

**Choosing migration rates.** Mutant fixation in the fastest deme is at the root of the impact of population structure on resistance evolution that we evidenced here, in particular in Fig 2. Suppose that a successful R mutant appears in the fastest deme: S individuals from connected demes can migrate to the fastest deme and take over there, which takes the population back to the initial fully S state. Thus, to make different structures as comparable as possible, we set migration rates so that the rate of invasion of one deme by the connected demes is the same across structures.

For the square lattice or grid, this leads to $\gamma_G = \gamma(D-1)/4$, where $\gamma$ is the per capita migration rate in the clique. Note that this condition also yields an equal total exchange rate in the clique and the lattice.

For the star, we first remark that a successful mutant is $(D-1)$ times more likely to first appear in a leaf than in the center. Neglecting the case where a successful mutant first appears in the center leads to the condition $\gamma_O = (D-1)\gamma$. Note that this condition does not coincide with setting equal overall exchanges between demes in the star and in the clique.

Finally, for the line, we note that for $D \gg 1$, successful mutants are much more likely to appear in a deme that is not at an end of the line. We thus choose migration rates such that the rate of invasion of one non-end deme by neighboring ones in the line is the same as in the clique: $\gamma_L = \gamma(D-1)/(\alpha+1)$, and $\gamma_R = \alpha\gamma_L$. Note that this condition slightly differs from setting equal overall exchanges in the line and in the clique, which would give $D$ instead of $D-1$ in the numerator of the expression of $\gamma_L$.

**Simulation results.** Fig S6C shows the survival probability of bacterial populations with different spatial structures with $D = 16$ demes. We observe that the fully subdivided population with $\gamma = 0$ has

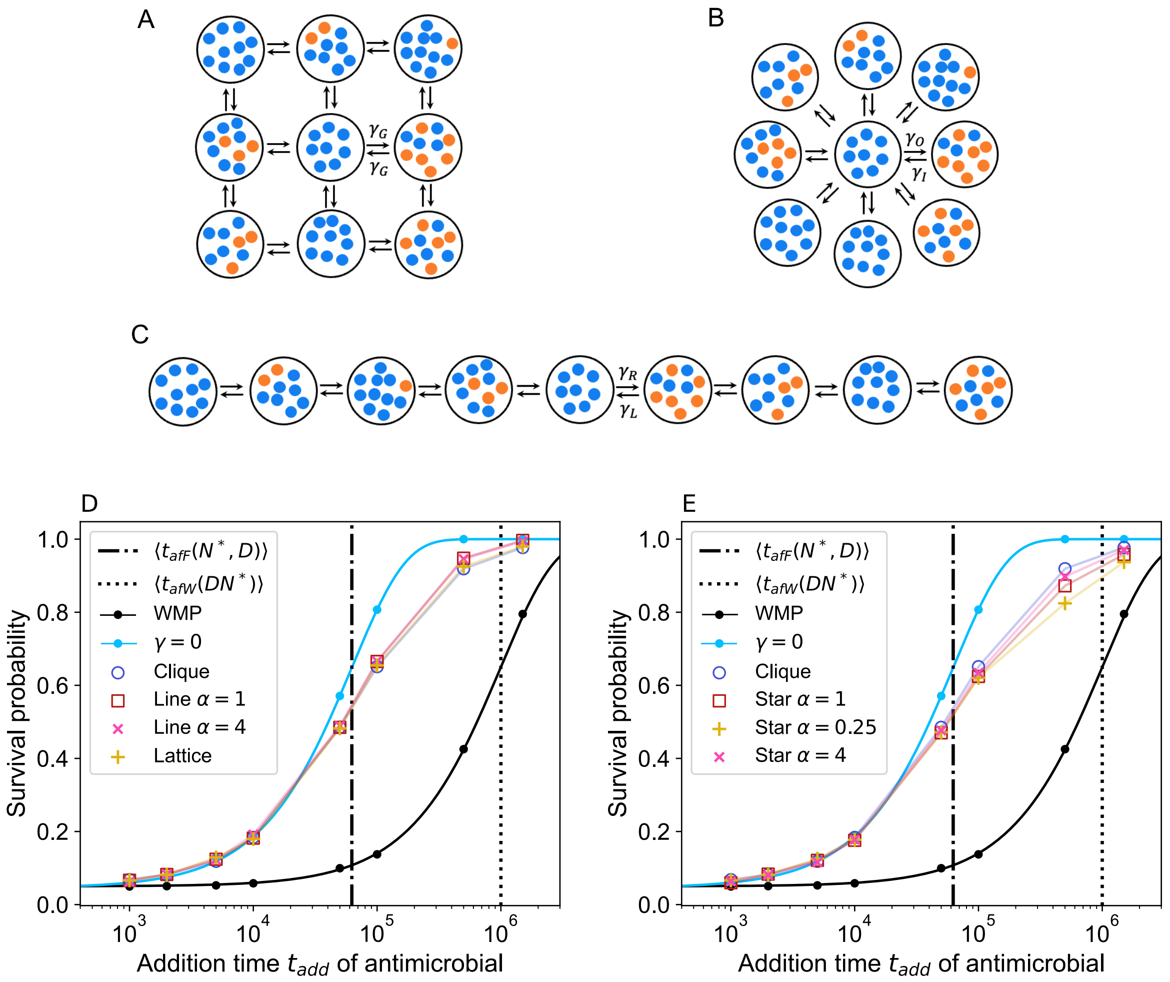

Figure S6: **Survival probability of a bacterial population to biostatic drug, for different spatial structures.** Panels A, B, C: Schematics of the spatial structures considered, shown with $D = 9$ demes for visualization ease: a square lattice with migrations $\gamma_G$ to each nearest neighbor (A), a star with and migration rates $\gamma_O$ from the center to a leaf and $\gamma_I = \alpha\gamma_O$ from a leaf to the center (B), and a line with migration rates $\gamma_L$ to the left and $\gamma_R = \alpha\gamma_L$ to the right (C). Panels D, E: Survival probability of a population starting from a sensitive inoculum versus drug addition time $t_{\text{add}}$, for different spatial structures: line with different migration asymmetries and lattice (D), star with different migration asymmetries (E). Results for a fully subdivided population with no migrations ("$\gamma = 0$"), a clique, and a well-mixed population are shown in both panels for reference. For the fully subdivided and well-mixed populations, the prediction from Eq 4 is shown as a solid line. Other lines are guides for the eye. Parameter values: $D = 16$, $K = 100$, $f_R = 1$ (no resistance cost), $f_S = 1$ before drug addition, $f_S = 0$ when antibiotic is added, $g = 0.1$, $\mu = 10^{-5}$, $\gamma = 10^{-6}$ for the clique, $\gamma_G = 3.75 \times 10^{-6}$ for the lattice, $\gamma_O = 15 \times 10^{-6}$ and $\gamma_I = \alpha\gamma_O$ for the star, $\gamma_L = \gamma_R = 7.5 \times 10^{-6}$ for the line with $\alpha = 1$, $\gamma_L = 1.2 \times 10^{-5}$ and $\gamma_R = \alpha\gamma_L$ for the line with $\alpha = 4$. Each result is obtained from $10^4$ simulation replicates.

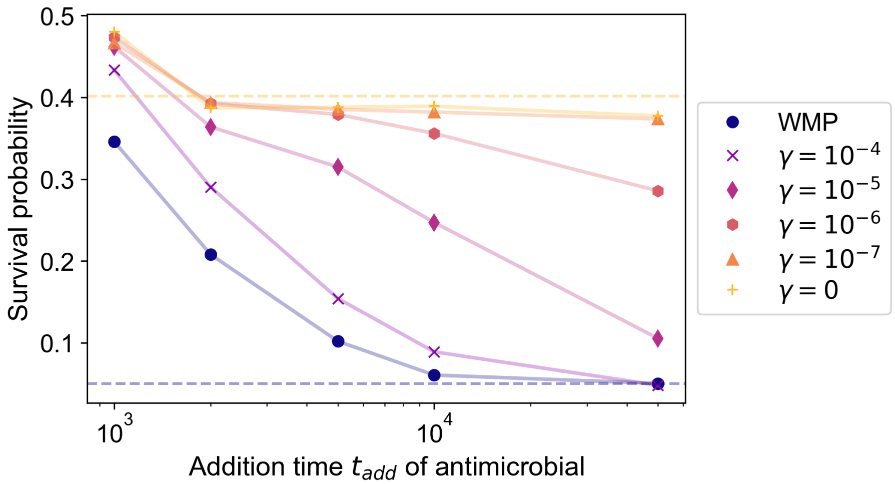

Figure S7: **Survival probability of a structured bacterial population with mutants in the inoculum upon addition of a biostatic drug.** The survival probability is plotted versus the addition time $t_{\text{add}}$ of antimicrobial. We consider a clique population structure with various migration rates. Results for a well-mixed population with same total size ("WMP") are presented as reference. Horizontal dashed lines: analytical predictions of the probabilities to still have mutants in the system in the long term (dark blue: well-mixed population; yellow: fully subdivided population with $\gamma = 0$). Data is obtained from $10^4$ replicate simulations in each case. Parameter values: $K = 100$, $D = 10$, $f_S = 1$ without drug, $f_S = 0$ with drug, $f_R = 1$, $g = 0.1$, $\mu = 0$. Initial percentage of mutants in each deme (and in the well-mixed population): 5%.

### 6 Concrete examples of spatially structured populations

In hosts, different organs can feature bacterial colonization and possess spatial structures with numerous demes. Here, we provide some details on the orders of magnitudes mentioned in the Discussion.

**Estimate of the number of intestinal crypts.** Bacteria are often found in intestinal crypts, which are glandular structures located at the base of the intestinal lining. In mice, intestinal crypts are densely packed, with approximately $10^5$ crypts in the intestine. This estimate arises from the typical distance of 30 µm between crypts, an intestinal length of 8 cm, and a circumference of 9 mm [13]. Besides, an analysis of 8 µm-thick slices of mouse colon crypts revealed the presence of 15 to 35 bacteria in each slice [14]. Considering a crypt depth of approximately 100 µm [15], this yields a range of 150 to 450 bacteria per crypt. For each crypt, the bacterial population is estimated to range from 100 to 400 bacteria [14]. Note that a given infection may not affect all crypts in a host.

**Estimate of the number of skin pores.** Bacteria also colonize skin pores in humans [16]. In the skin of the human face and nose, there are approximately 20 to 30 pores per $0.8\,\mathrm{cm}^2$ [17]. With the average area of the face being around $600\,\mathrm{cm}^2$, this corresponds to roughly $2 \times 10^4$ pores.
